# No effect of education on Telomere Length: a natural experiment in aging individuals

**DOI:** 10.1101/2025.01.17.633604

**Authors:** Nicholas Judd, Rogier Kievit

## Abstract

Telomere length is increasingly used as a proxy for an ‘aging exposome’ – an index of non-genetic environmental (and behavioral) exposures that an individual encounters, leading to differences in rates of aging. One of the largest associations with telomere length is schooling, leading to the suggestion of long-lasting protective effects of (additional) education. However, for ethical and practical reasons, education cannot generally be randomized, precluding an understanding of the causal effect of education on health outcomes. Here, we leverage a well-established policy change in the UK to test whether an additional year of education caused differences in telomere length decades later. Employing a fuzzy regression discontinuity design, we find no effect of education on telomere length in the UK BioBank across multiple analytic frameworks (continuity, local randomization & Bayesian approaches) and a series of robustness tests, such as increasing the analysis window around the cutoff, and changing the prior, similarly find no effect of an additional year of education on telomere length. This robust finding contrasts the prevailing ‘aging exposome account’ of telomere length and offers a cautionary note in attributing causal power to environmental factors based on observational differences.

## Introduction

Telomeres are the protective caps of our chromosomes – repeating nucleotide sequences – shortening with each cell cycle as we age. Rare inherited telomere syndromes in humans and experimental animal work provide causal evidence for a link between shorter telomeres and senescence (Blackburn *et al*., 2015). Despite telomere length’s high heritability (∼70%) (Broer *et al*., 2013), there is evidence that it can be affected by non-genetic factors (Epel *et al*., 2004; Adler *et al*., 2013; Bountziouka *et al*., 2022). These findings have led to prominent accounts of telomere length as a proxy for *‘the aging-exposome’* – an index of non-genetic environmental (and behavioral) exposures that an individual encounters, leading to differences in rates of aging (Samani & van der Harst, 2008; Blackburn *et al*., 2015; Rentscher *et al*., 2020).

A large body of observational work has followed from this logic – linking telomere length with important individual characteristics thought to alter the pace of aging and disease onset (e.g., education, stress, and socioeconomic status). In general, healthy behaviors and positive exposures are associated with longer telomeres, whereas negative environmental influences are associated with shorter telomeres (Rentscher *et al*., 2020; Bountziouka *et al*., 2022). However, the causal involvement of these modifiable environmental factors on telomere length has been repeatedly questioned (Simons, 2015; Notterman & Schneper, 2020; Rentscher *et al*., 2020).

Education has been singled out as a particularly important environmental influence on health outcomes and telomere length (Adler *et al*., 2013; Bountziouka *et al*., 2022). Yet, a range of societal and inter-individual characteristics lead some individuals to attain more education than others, making it challenging to disentangle the (potential) causal impact of education on telomere length (Pepper & Nettle, 2017). Moreover, evidence suggests that genetic differences may masquerade as environmental influences (via gene-environment correlations), further complicating causal and mechanistic inferences (Paige Harden, 2021).

One solution to this challenge is the natural experimental design – offering an avenue to study *causation* in phenomena that cannot be randomized for practical or ethical constraints (Cattaneo & Titiunik, 2022). For instance, policy changes have been essential in assessing causality between education and health outcomes (Cattaneo & Titiunik, 2022). Natural experimental designs offer a powerful paradigm to test if lifestyle factors can modify telomere length. However, despite explicit calls (Rentscher *et al*., 2020) for these designs to be used to address such challenges, they remain severely underutilized. Here, we demonstrate their value – by isolating the causal effect of additional education on telomeres with a natural experiment.

On September 1st 1972, the minimum mandatory age to leave school was increased from 15 to 16 years of age in England, Scotland, and Wales with the Raising of School Leaving Age Order (called here ROSLA) (*The Raising of the School Leaving Age Order*, 1972; Davies *et al*., 2018; Judd & Kievit, 2024a). In line with prior associational work, we expect an additional year of education from this policy change to increase telomere length. To test this, we pair cutting-edge causal inference methods from econometrics with the UK Biobank – a cohort study that overlaps with ROSLA and has leukocyte telomere length measured in 474,074 participants (Davies *et al*., 2018; Codd *et al*., 2022).

## Results & Discussion

The policy change ROSLA is a well-validated natural experiment (Clark & Royer, 2013; Barcellos *et al*., 2018, 2023; Davies *et al*., 2018; van de Weijer *et al*., 2023; Judd & Kievit, 2024b) – shown to increase qualifications, income, and cognition while also decreasing adverse health outcomes such as diabetes and mortality. Education is consistently highlighted as a major factor underlying heterogeneous neurodevelopment thought to contribute to a ‘cognitive reserve’ – buffering against adverse aging effects and is one of the largest modifiable risk factors associated with telomere length (Adler *et al*., 2013; Bountziouka *et al*., 2022).

First, we replicated prior work (Adler *et al*., 2013; Blackburn *et al*., 2015; Bountziouka *et al*., 2022; Codd *et al*., 2022), finding telomere length to be significantly *negatively* associated with chronological age (B = -.021, CI = [-.021,-.022], p < .001, β = -.123) and *positively* associated with years of education (B = .020, CI = [.019,.022], p < .001, β = .054) in 256,243 individuals (Appendix 1). Each additional year of education corresponds to one less year of chronological aging on telomere length (Figure 1a). This leads to our core hypothesis: does education *causally increase* telomere length, thereby buffering the adverse consequences of natural aging?

**Figure 1.**
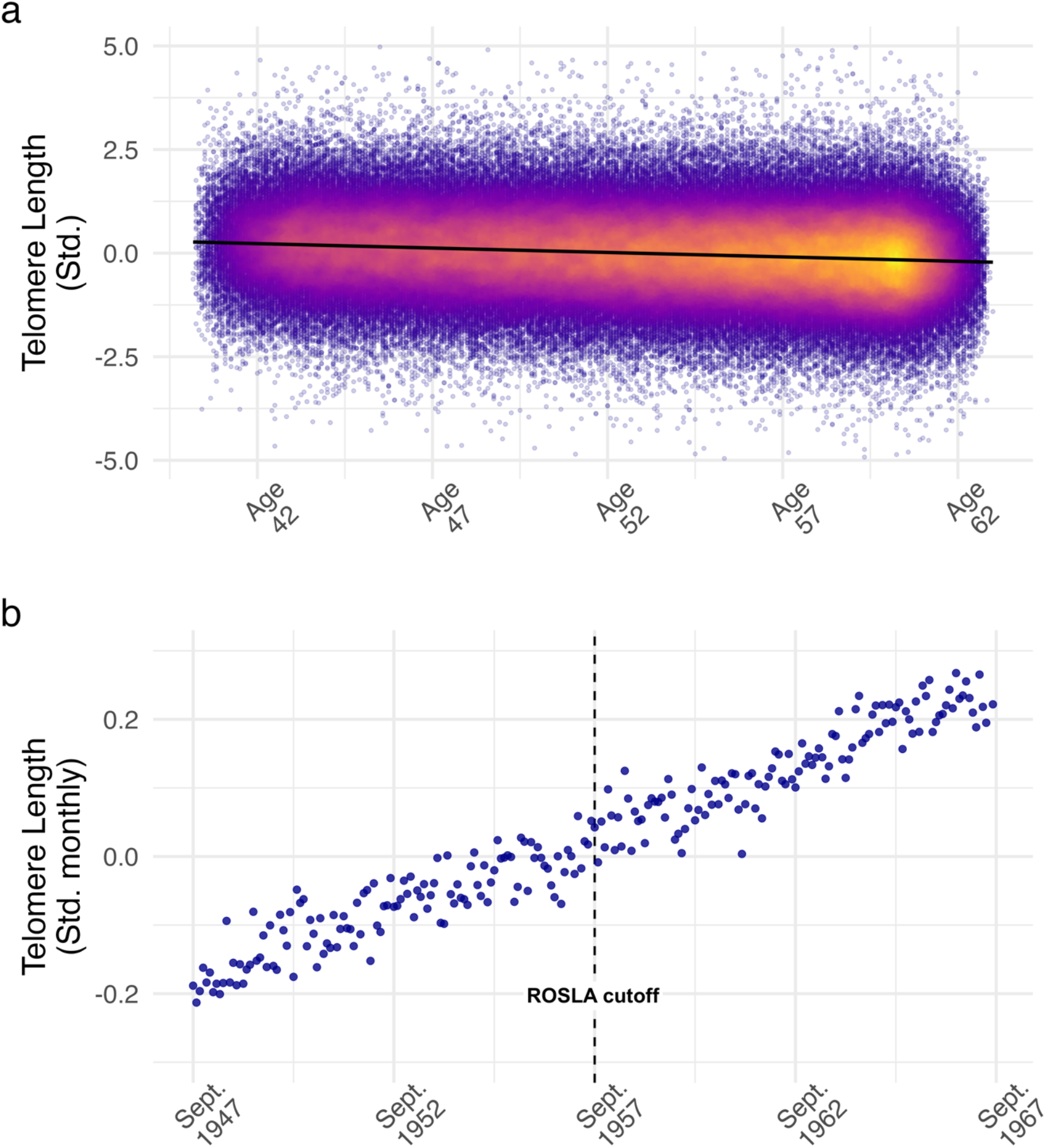
Illustrates the relationship of telomere length (from leukocyte cells) with age (*p* < .001) and an additional year of education (*p* > .05 across a series of tests; Sup Table 2 & 3) from the Raising of School Leaving Act (ROSLA). **Panel *a)*** gives an overview of our sample (N = 256,243) with a density scatter plot showing the well-established relationship of telomere length with age. **Panel *b)*** is a regression discontinuity plot of monthly averaged Telomere Length (from leukocyte cells) plotted by the participant’s date of birth in months (our running variable). Each dot reflects the average value for individuals born in that month. The dashed line corresponds to Sept. 1957 the date of birth inclusion cutoff for an additional year of mandatory education from the policy change ROSLA. We found no effect (p >.05, BF_01_ > 20, ROPE_0.1SD_ > 95%) from an additional year of education (ROSLA) on telomere length illustrated by a continuous line around the cutoff and tested via a series of fuzzy regression discontinuity analyses (Sup Table 2, 3 & 4).

Pairing the UK Biobank with an educational policy change (ROSLA) allows us to isolate the effect of an additional year of education in a large sample. Children were allocated an additional education based on their date of birth, creating an arbitrary cutoff whereby individuals near the cutoff are similar in many respects (Barcellos *et al*., 2018; Davies *et al*., 2018; Judd & Kievit, 2024a) except for the extra year of mandatory schooling. In our sample, we found ROSLA adherence to be almost 100%, with a first-stage estimate of 16% (Sup. Figure 4a & Sup. Table 4) – that is, 16% of adolescents who would have left school before ROSLA were now forced to stay in school an additional year.

Using fuzzy regression discontinuity methods (Cattaneo *et al*., 2017; Cattaneo & Titiunik, 2022), we compare the telomere length of individuals *immediately* affected by the policy change (Sept. 1957; N = 961) to those born one month earlier and unaffected (August 1957; N =1016). We find no effect (*p* > .05) of an additional year of education on telomere length (Figure 2; Sup Table. 2). In a Bayesian approach, with the same individuals, we find ‘very strong’ evidence (BF_01_ = 43.48) in favor of the null hypothesis for ‘no effect from education on telomere length’ (Sup. Appendix 3). A series of robustness tests, such as increasing the analysis window around the cutoff (Sup. Table 2), changing the prior (Sup. Table 3), using region of practical equivalence boundaries, and continuity-based approaches (Sup. Appendix 4), similarly find no effect of an additional year of education on telomere length.

**Figure 2.**
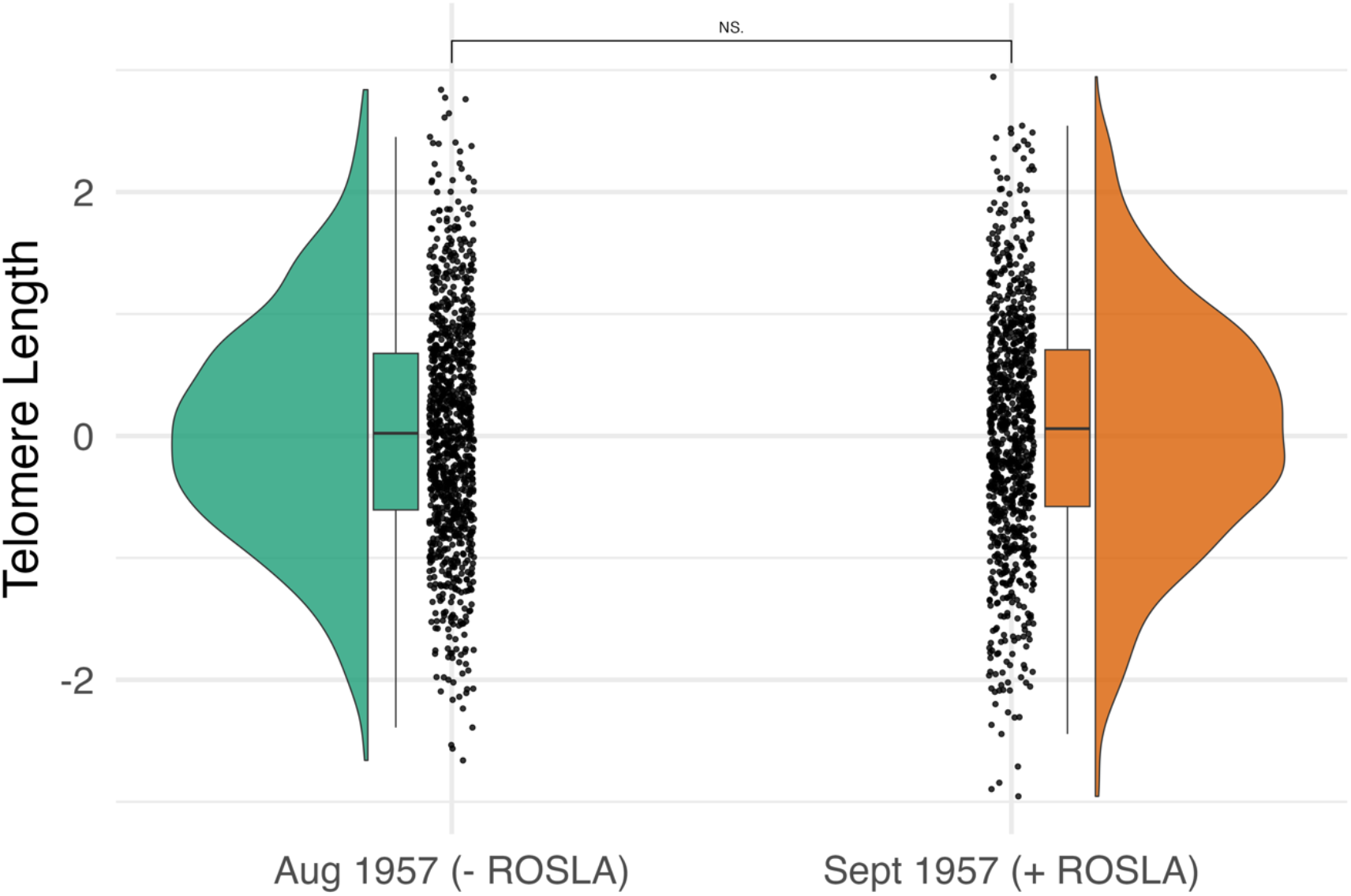
A raincloud plot illustrating telomere length for individuals born in August 1957 (N=1016; who were unaffected by the policy change ROSLA) and individuals born in September 1957 (N=961; who had to complete an additional year of mandatory education from the 1973 ROSLA). A local randomization analysis found no significant differences (*p* > .05) in telomere length between these two groups. A Bayesian approach provided evidence in support of the null hypothesis using Bayes Factors (*BF*_*01*_ *= 43*.48) and ROPE’s (100% of the 95-HDI for an effect size of .1SD; 77% for an effect size of .05SD). This null result was robust to increasing the analysis window (Sup. Table 2), Bayesian prior sensitivity testing (Sup. Table 3), and continuity regression discontinuity methods (Sup. Table 4).

In summary, we replicate the *associational* effect of telomere length on education, finding each additional year of education to be a similar size as one less year of chronological aging (Bountziouka *et al*., 2022). However, we did not find a *causal* effect of education on telomere length, instead, providing very strong evidence in favor of the null hypothesis. Our analyses are robust to analytical choices, observed in a large sample, and employ a well-validated natural experimental design.

Prior work – in the same sample with identical quantitative approaches – has found this additional year of education to impact cognitive, behavioral, and health outcomes (Barcellos *et al*., 2018, 2023; Davies *et al*., 2018) demonstrating the ROSLA policy change had causal impacts on some outcomes (although see (Clark & Royer, 2013; van de Weijer *et al*., 2023; Judd & Kievit, 2024a)). A major hurdle in this field is the degree of analytic flexibility; for instance, two recent papers using ROSLA studying the same outcome, in the same sample (dementia risk), found opposite conclusions (Barcellos *et al*., 2025; Monnet *et al*., 2026). Preregistration, robustness checking, and automatic bandwidth selection can help standardize regression discontinuity methods, yet a lack of suitable continuity-based techniques for discrete running variables still limits the field (Cattaneo *et al*., 2023).

One limitation on generalization is that the ROSLA intervention only increased education between the ages of 15 and 16 – changing the timing or duration of the intervention could affect biological aging markers such as telomere length yet this remains to be empirically supported. In addition, UK Biobank respondents are more educated than the general population, indicative of *‘healthy volunteer bias’* (Fry *et al*., 2017; van Alten *et al*., 2024). For this reason, the percentage of adolescents impacted by ROSLA is lower than previously reported (25% vs 16%; SI Table 4)(Clark & Royer, 2013), limiting our power across some specifications (Sup. Table 2). Our findings are in agreement with prior Mendelian randomization work, concluding if there is an effect from education on telomere length it is not of clinical significance (Bountziouka *et al*., 2022).

An additional year of education is a substantial intervention. As such, our robust null effect challenges an ‘aging exposome account’ of telomeres and adds empirical weight to the repeated warnings (Simons, 2015; Notterman & Schneper, 2020; Rentscher *et al*., 2020) that the *causal* nature of environmental exposures on telomere length in humans is yet to be established. Since the vast majority of prior telomere work in humans is observational, our findings suggest that caution is warranted if such previous findings are interpreted as treatment- or intervention targets in the absence of evidence to warrant causal claims.

## Code and Data Availability

All code is publicly available (https://github.com/njudd/EduTelomere). The data is also publicly available yet must be accessed via the centralized UK BioBank repository (https://www.ukbiobank.ac.uk). This research has been conducted using the UK Biobank resource under application number 23668.

## Funding & Acknowledgments

Nicholas Judd is supported with eScience and Jacobs fellowships. Rogier Kievit is supported by a Hypatia fellowship from the RadboudUMC. This research has been conducted using the UK Biobank resource under application number 23668. We would like to thank Barbara Sakic, Barbara Franke, Andre Marquand, Jan Buitelaar, Nina Roth Mota, Janita Bralten, and Ward de Witte for help in navigating and accessing the UK Biobank.

## Ethical Statement

UK Biobank has ethical approval from the North West Multi-centre Research Ethics Committee (MREC) as a Research Tissue Bank (RTB) (approval number: 11/NW/0382). This consortium has its own independent ethics advisory committee (https://www.ukbiobank.ac.uk/learn-more-about-uk-biobank/governance/ethics-advisory-committee), that assures the UK Biobank abiders by the ethics and governance framework (https://www.ukbiobank.ac.uk/media/0xsbmfmw/egf.pdf). Written informed consent was obtained from all participants.

## Supplementary Information

### Sup. Appendix 1: Descriptive and Sample Characteristics

Using the UK BioBank ^1^ we included participants 10 years on either side of the ROSLA date of birth cut-off (Sept 1947 – August 1967) who reported being born in England, Wales, or Scotland and had leukocyte telomere length (*cf*.^*2*^; field 22191). We logged and standardized (mean = 0, SD = 1) telomere length as recommended^2^.

Our running variable was an individual’s date of birth in months, which was centered so zero corresponded to the start of ROSLA (September 1957). Age is measured in months, which is the time between a participant’s DOB and visit to a UK Biobank center. Age and the running variable encode similar information (R = .99).

Education was defined as the age at which a participant left full-time education, those completing college were not asked this question and therefore recoded to 21 years as is common in the literature^3-5^. Our first-stage ‘fuzzy’ outcome was the education variable dummy coded whether a participant left school at 16 or above (EduAge16). The R language (v 4.4.0) was used for all analyses (https://github.com/njudd/EduTelomere).

Due to extensive prior work on ROSLA^6,7^ and in this sample specifically ^3,4,8-10^, including our own work ^5^, we did not conduct classic RD design falsification tests (density test, predetermined covariates, placebo outcomes). Since we did not find an effect of education, we did not conduct placebo outcome tests.

**Sup. Table 1:** Descriptive Table. Descriptive table of included variables: leukocyte telomere length (LTL); outlier removed LTL (LTL_3SD); age measured in months divided by 12 (age); running variable of date of birth in months when 0 is September 1957 the ROSLA birthdate cutoff (running_var); running_var standardized (running_var.s); age left full-time education (EduAge); EduAge standardized (EduAge.s); a dummy coded variable for participants who left full-time education 16yrs or after (EduAge16); sex.

| Variable | N | Mean | SD | Median | MAD | Min | Max |
| --- | --- | --- | --- | --- | --- | --- | --- |
| LTL | 256243 | 0.00 | 1.00 | 0.00 | 0.95 | -11.02 | 12.25 |
| LTL_3SD | 254816 | 0.00 | 0.96 | 0.00 | 0.95 | -3.00 | 3.00 |
| age<sup>†</sup> | 256243 | 52.74 | 5.86 | 53.33 | 7.29 | 40.17 | 63.00 |
| running_var<sup>†</sup> | 256243 | -16.68 | 69.43 | -24.00 | 87.47 | -120.00 | 119.00 |
| running_var.s<sup>†</sup> | 256243 | 0.00 | 1.00 | -0.11 | 1.26 | -1.49 | 1.95 |
| EduAge | 256243 | 18.36 | 2.64 | 18.00 | 4.45 | 5.00 | 35.00 |
| EduAge.s | 256243 | 0.00 | 1.00 | -0.14 | 1.68 | -5.06 | 6.30 |
| EduAge16 | 35095 |  |  |  |  |  |  |
| (FALSE) | (13.7%) |  |  |  |  |  |  |
| EduAge16 | 221148 |  |  |  |  |  |  |
| (TRUE) | (86.3%) |  |  |  |  |  |  |
| Sex | 141536 |  |  |  |  |  |  |
| (Female) | (55.2%) |  |  |  |  |  |  |
| Sex | 114707 |  |  |  |  |  |  |
| (Male) | (44.8%) |  |  |  |  |  |  |

**Note:** <sup>†</sup> Age and the Running Variable encode the same information (Pearson $R = .99$ )

**Sup. Figure 1:**
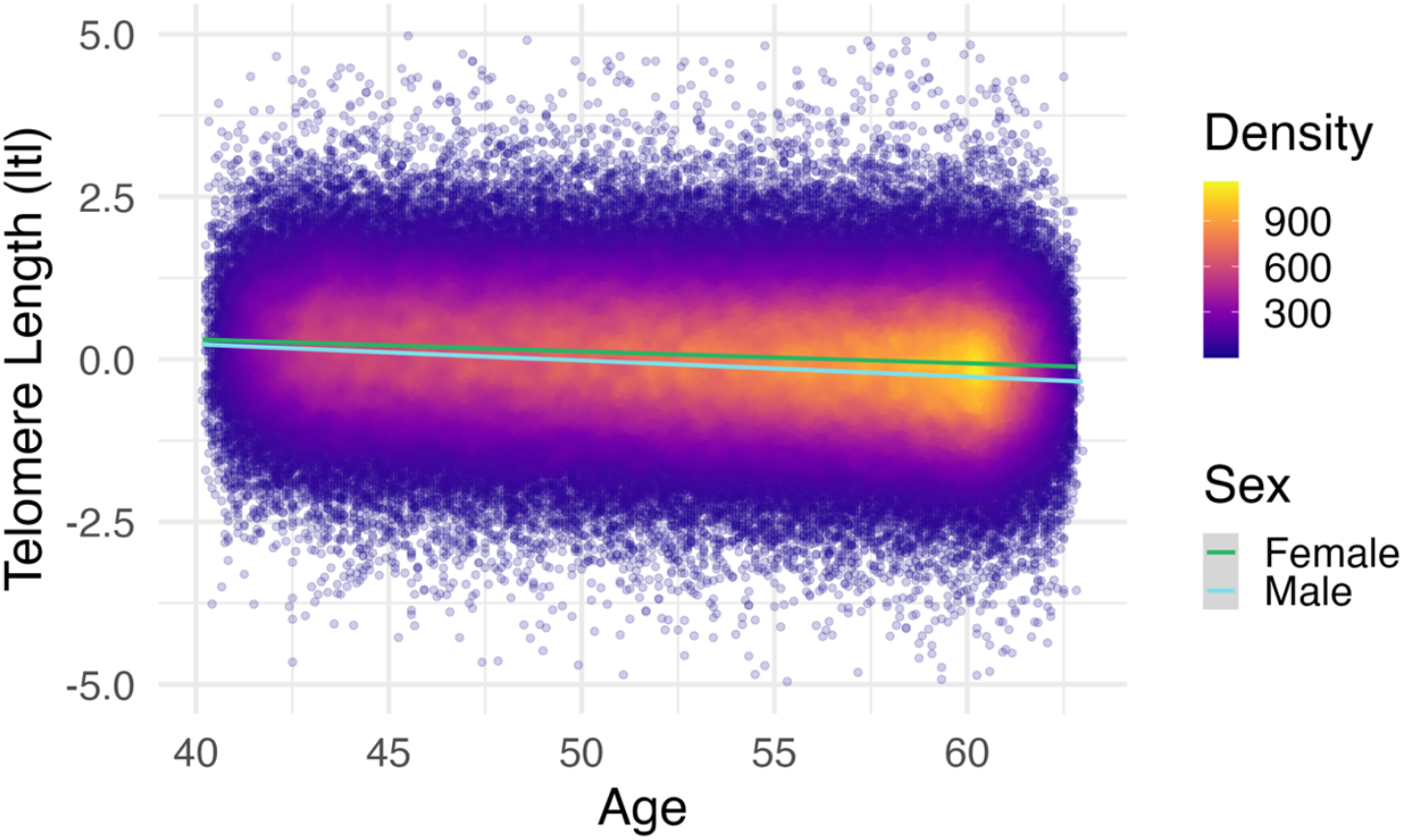
Density plot of Leukocyte Telomere Length by Age and Sex. A density plot with each participant’s value of telomere length (y-axis) plotted individually (N = 256,144) by age (x-axis). Linear trends are plotted by sex (Female & Male). The y-axis was limited to 5 standard deviations, this excluded 99 participants (∼.0004%).

### Sup. Appendix 2: Fuzzy local-randomization regression discontinuity

Local-randomization regression discontinuity analysis is the natural framework ^11,12^ to test the efficacy of an additional year of education from ROSLA on telomere length as our running variable was discrete (e.g., date of birth in months). We used the smallest window possible by comparing individuals born in August 1957 (non-affected) to individuals born in September 1957 (ROSLA-affected).

To implement frequentist randomization inference, we used ‘rdrandinf’ function from the rdlocrand package (v 1.0) with the default options (e.g., uniform kernel, p = 0) while setting ‘wmasspoints’ to TRUE. As mentioned above, our fuzzy outcome was a true/false vector on whether the individual completed 16 years of education (EduAge16). We report large sample p-values (Sup. Table 2) for both intent-to-treat (ITT) and two-stage least squares (TSLS) estimates. We report the minimal effect size possible to detect with at least 80% power. These power analyses are complemented by Bayes Factors and ROPE boundaries in our Bayesian analysis (Sup. Table 3), offering converging evidence in support of the null hypothesis. Lastly, as a robustness test, we expanded the analysis window to five months on either side of the cutoff.

**Sup. Table 2:** Leukocyte Telomere Length Results (fuzzy local-randomization RD) Supplementary results of fuzzy local-randomization regression discontinuity (RD) analyses using the rdlocrand package (v 1.0) for a 1-month window (1-month w) and a 5-month window (5-month w). We report a large sample p-value. (l) stands for the left of the ROSLA cutoff (not affected), while (r) stands for the right of the cutoff; therefore, individuals affected by the policy change ROSLA. We report both Intention to treat (ITT) and two-stage least square estimates (TSLS). See Sup. Appendix 3 for Bayesian null hypothesis testing.

| Statistic | N (l) | Mean (l) | N (r) | Mean (r) | Estimate | *p* | >80% power |
| --- | --- | --- | --- | --- | --- | --- | --- |
| 1-month window analysis: |  |  |  |  |  |  |  |
| ITT | 1016 | .052 | 961 | .042 | -.010 | .838 | .13 SD |
| TSLS | 1016 | .052 | 961 | .042 | -.044 | .831 | .57 SD |
| 5-month window analysis: |  |  |  |  |  |  |  |
| ITT | 5295 | .027 | 4889 | .039 | .013 | .528 | .06 SD |
| TSLS | 5295 | .027 | 4889 | .039 | .081 | .519 | .35 SD |

**Sup. Figure 2:**
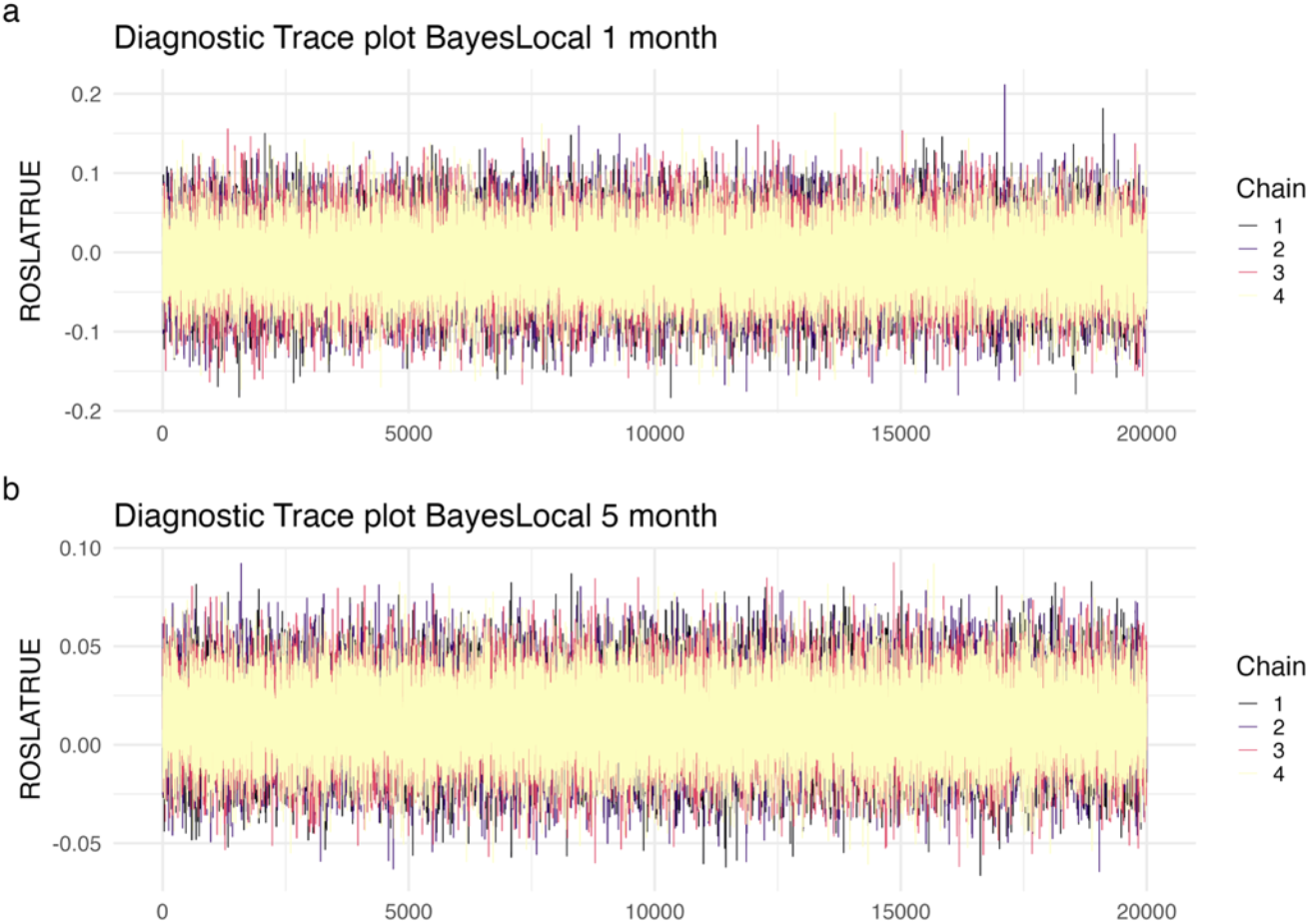
Bayesian diagnostic trace plot (prior 1,0)

**Sup. Figure 3:**
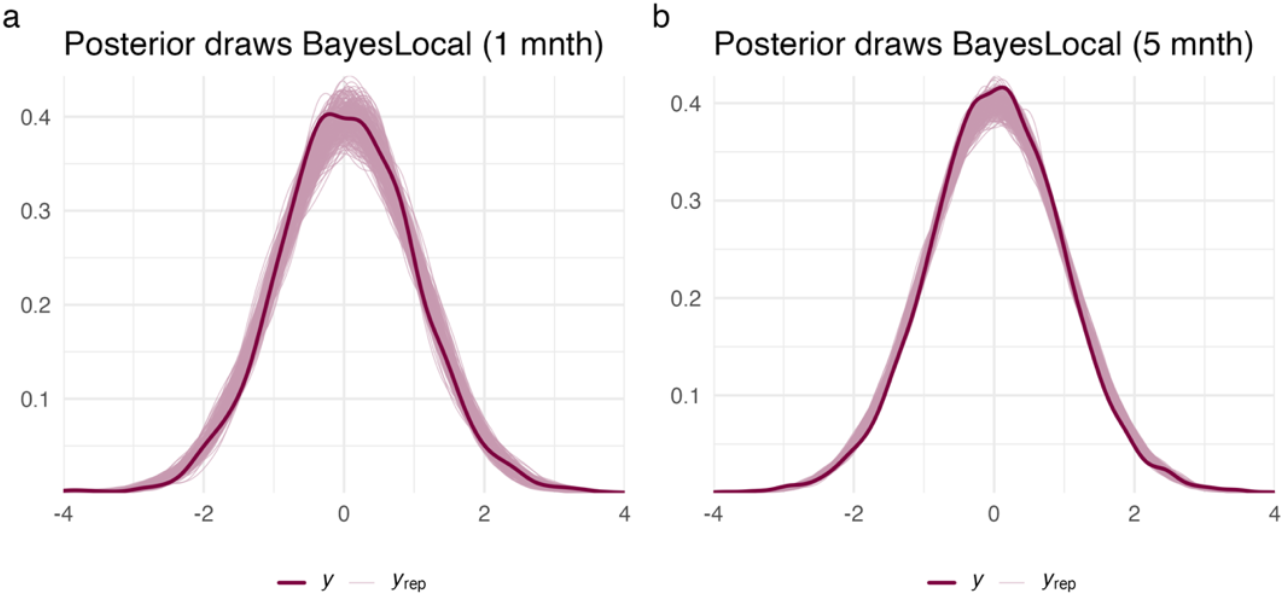
Bayesian diagnostic posterior predictive check plot (prior 1,0)

### Sup. Appendix 3: Bayesian local-randomization analysis

We conducted a Bayesian local-randomization analysis as an additional robustness test. Local randomization regression discontinuity assumes a small window around the cut-off where participants are ‘as if random’ ^12,13^. Choosing the optimal window becomes a bias versus variance tradeoff – as the window increases (around the cutoff) variance decreases since more participants are included, yet bias also increases as the included participants are further away from the cutoff and, in turn, less similar. For this reason, we chose the smallest window possible with our discrete running variable, comparing participants in (*ω* = 1 month; August vs September 1957), then expanded to a 5-month window around September 1^st^, 1957 ^5^. We then dummy-coded whether the participant was impacted by the natural experiment ROSLA, and tested this term on standardized log transformed telomere length (as outlined in Appendix 1).

Models were fit in R (v. 4.4.0) with rstanarm ^14^ with Markov Chain Monte Carlo sampling of 80,000 iterations over 4 chains. Our priors used a normal distribution and were centered at 0. We sensitivity-tested our prior for the effect from ROSLA by changing the standard deviation to three values (0.5, 1, 1.5). Model diagnostics were checked with trace plots, posterior distributions, and rhat values (<1.05) and found acceptable (Figure 3 & 4) ^15^. We then computed Bayes Factors using the bayestestR package ^16^ using the Savage-Dickey density ratio with a point-null of zero. The strength of the evidence was interpreted according to Jeffrey’s criteria ^17^. We report results from the normal prior with a mean of zero and a standard deviation of one.

Regions of practical equivalence, or ROPEs, are an alternative Bayesian null hypothesis testing framework whereby a region *or* minimal effect size of interest is set. We used the ROPE function from the bayestestR package for regions of 0.1 SD and 0.05 SD – this function uses the 95% HDI of the posterior to calculate ROPEs. The percentage reflects the amount of the posterior (95% HDI) inside a ‘region of practical equivalence’ if 95% (or another arbitrary alpha) lies within this region, the null hypothesis is confirmed. Alternatively, if an effect is present, it should be fully (to X% of alpha) outside this region. If the posterior is neither outside nor instead of the ROPE then it is not informative for the alternative or null hypothesis for the tested effect size (in our case 0.1 SD and 0.05 SD).

**Sup. Table 3:** Leukocyte Telomere Length Results (Bayesian local-randomization) Supplementary results of our Bayesian local-randomization analyses for a 1-month window (A) and a 5-month window (B) across three normally distributed priors. We report the prior with a mean of zero and an SD of 1.

| Model | N | Posterior mean[sd] | 95% HDI | BF<sub>01</sub> | ROPE .1 SD | ROPE .05 SD |
| --- | --- | --- | --- | --- | --- | --- |
| **A. 1-month window** |  |  |  |  |  |  |
| *Normal(0, 0.5)* | 1977 | -.009[.044] | (-.096, .078) | 21.7 | 100% | 77% |
| *Normal(0, 1)* | 1977 | -.009[.044] | (-.096, .078) | 43.5 | 100% | 77% |
| *Normal(0, 1.5)* | 1977 | -.009[.044] | (-.096, .078) | 66.7 | 100% | 77% |
| **B. 5-month window** |  |  |  |  |  |  |
| *Normal(0, 0.5)* | 10184 | .013[.02] | (-.026, .051) | 41.7 | 100% | 99.6% |
| *Normal(0, 1)* | 10184 | .013[.02] | (-.026, .051) | 83.3 | 100% | 99.6% |
| *Normal(0, 1.5)* | 10184 | .013[.02] | (-.026, .051) | 125 | 100% | 99.6% |

### Sup. Appendix 4: Fuzzy local-linear regression discontinuity

We fit fuzzy local-linear regression discontinuity using the RDHonest package (v 1.0; https://github.com/kolesarm/RDHonest)^18^ using the default options of RDHonest (MSE-derived bandwidth estimation, triangular kernels, Holder class, local linear) due to its ability to optimally handle discrete running variables ^19^ (in our case, date of birth in months; DOB). We only included participants 10 years on either side of the ROSLA date of birth cut-off (Sept 1947 – August 1967) who reported being born in England, Wales, or Scotland and had telomere length (Outline in Appendix 1).

Due to extensive prior work on ROSLA^6,7^ in this sample ^3,4,8-10^, including our own work ^5^, we did not conduct classic RD design falsification tests (density test, predetermined covariates, placebo outcomes). Since we did not find an effect of education, we did not conduct placebo outcome tests.

To test the robustness of our finding, we re-ran the same model with two sensitivity tests: *1)* participants with more than 3 standard deviations in leukocyte telomere length removed (∼0.5% of the total data) and *2)* correcting for visit date measured in days from the first participant quadratically to increase precision. All models were also tested with Eicker-Huber-White clustered standard errors with UK Biobank site. We tested both visit date and visit date^^2^ as placebo outcomes, finding them *unrelated* (*p* < .05) to ROSLA. Placebo outcomes are a validity test of natural experiments. If a natural experiment is well-designed, placebo outcomes should be unrelated to the natural experiment (in this case, ROSLA). Our placebo outcome findings add to the extensive prior work on ROSLA in the UK Biobank ^3,5,8-10^, further justifying the use of ROSLA to study the effect of education.

**Sup. Table 4:** Leukocyte Telomere Length Results (fuzzy local-linear RD) Supplementary results of fuzzy local-linear regression discontinuity (RD) analyses for leukocyte telomere length using the A) main model, B) outlier corrected, and C) covariate corrected. Standard errors clustered by UK Biobank site with Eicker-Huber-White (EHW) estimation are also provided.

| Model | 1<sup>st</sup> stage | Eff. objs | Estimate | SE | 95% CI | p-value |
| --- | --- | --- | --- | --- | --- | --- |
| **A. Main model** |  |  |  |  |  |  |
|  | 0.16 | 37902.73 | 0.04 | 0.13 | -0.24, 0.31 | 0.79 |
| Cluster EHW | 0.16 | 37912.94 | 0.04 | 0.13 | -0.25, 0.32 | 0.80 |
| **B. Outlier model** |  |  |  |  |  |  |
|  | 0.16 | 36778.74 | -0.03 | 0.12 | -0.30, 0.24 | 0.84 |
| Cluster EHW | 0.16 | 36779.24 | -0.03 | 0.13 | -0.31, 0.26 | 0.85 |
| **C. Covariate model** |  |  |  |  |  |  |
|  | 0.16 | 39106.44 | 0.05 | 0.12 | -0.22, 0.32 | 0.74 |
| Cluster EHW | 0.16 | 39107.82 | 0.05 | 0.13 | -0.23, 0.32 | 0.75 |

**Sup. Figure 4:**
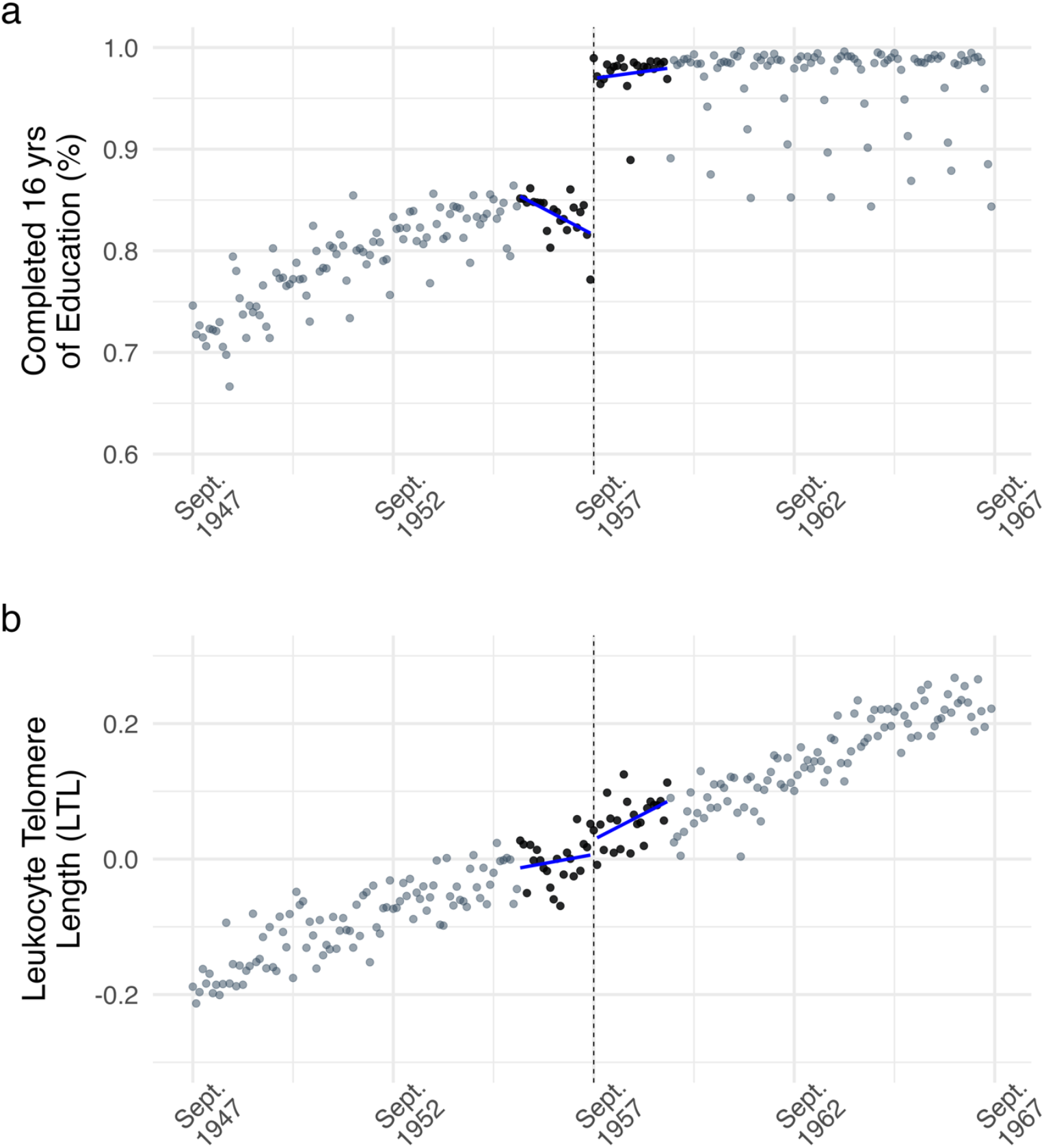
Fuzzy Local-linear Regression Discontinuity Plot. Regression discontinuity (RD) plot of monthly averaged ***a)*** number completing education until 16 years and ***b)*** Leukocyte Telomere Length (LTL) plotted by the participant’s date of birth in months (our running variable). Illustrating our first (a) and second (b) -stage outcomes from fuzzy local-linear regression discontinuity. Each dot reflects the average value for individuals born in that month, black dots represent the months used for inference (bandwidth = 22). The dashed line corresponds to Sept. 1957 the date of birth inclusion cutoff for an additional year of mandatory education from ROSLA. ***Note:*** *Examining panel a) shows some months with lower percentages of children staying in school until 16 years of age after ROSLA. These are summer-born children (June, July, August) who completed a similar number of years of education yet were able to leave at 15 (see* ^5^*). This pattern remained consistent on either side of ROSLA*.

